# Comprehensive intra-host infection kinetics reveals high arbo-flavivirus transmission potential by neglected vector species, *Aedes scutellaris*

**DOI:** 10.1101/2024.09.12.612593

**Authors:** Yudthana Samung, Jutharat Pengon, Chatpong Pethrak, Phonchanan Pakparnich, Saranya Thaiudomsup, Kittitat Suksirisawat, Anon Phayakkaphol, Songpol Eiamsam-ang, Thipruethai Phanitchat, Channarong Sartsanga, Tararat Jantra, Patchara Sriwichai, Natapong Jupatanakul

**Author notes:** **Corresponding authors:** Natapong Jupatanakul, Patchara Sriwichai. These authors contributed equally to this work (co-first authors) (Yudthana Samung, Jutharat Pengon, Chatpong Pethrak).

## Abstract

**Background:** Dengue virus (DENV) and Zika virus (ZIKV) are primarily transmitted by *Aedes* mosquitoes. As most studies on vector competence have focused on *Aedes aegypti* and *Aedes albopictus* while neglecting other *Aedes* species, it is possible that the transmission risks might be underestimated. it is necessary to examine additional species that could potentially serve as competent vectors. This is particularly important considering the potential expansion of their geographical range due to climate change or species-specific vector reduction interventions.

**Methodology/Principal Findings:** In this study, we examined the infection kinetics and transmission potential of *Aedes scutellaris* from Thailand, comparing to *Ae. aegypti* and *Ae. albopictus*. Our findings demonstrated that *Ae. scutellaris* and *Ae. albopictus* had lower rates of midgut infection compared to *Ae. aegypti* due to smaller blood meal sizes during feeding. However, once the infection has established *Ae. scutellaris* exhibited efficient replication of ZIKV and DENV1-4 in the midguts, secondary organs, and salivary glands. Notably, *Ae. scutellaris* had a low salivary gland escape barrier, with comparable transmissibility as *Ae. aegypti* when inoculated with the same viral load.

**Conclusion:** This study highlights the potential of *Ae. scutellaris* as a vector for DENV and ZIKV and emphasizes the importance of considering neglected mosquito species in arbovirus transmission and surveillance efforts.

**Author Summary:** Dengue and Zika are viral infections caused by arthropod-borne flaviviruses, and spread primarily through the bite of infected *Aedes* mosquitoes. Most research on DENV and ZIKV transmission has primarily focused on *Aedes aegypti* and *Aedes albopictus* while other *Aedes* species are overlooked, thus the epidemiology of the transmission might be underestimated. With climate change together with species-specific mosquito population reduction interventions these neglected *Aedes* species could become increasingly important in sustaining virus transmission. In this study, we examined *Aedes scutellaris*, a mosquito species that co-habitats with *Ae. aegypti* and *Ae. albopictus*, to assess its ability to transmit DENV and ZIKV using a combination of blood feeding and intrathoracic injection methods. Our findings show that although *Ae. scutellaris* had lower initial infection rates due to smaller blood meals, DENV and ZIKV were able to replicate and transmit at levels comparable to *Ae. aegypti* when exposed to similar virus loads. This highlights the need to study a broader range of species to improve virus control and outbreak prevention strategies.

## 1. Introduction

Dengue and Zika infections are both caused by arthropod-borne flaviviruses (arboflaviviruses) belonging to the Flaviviridae family. Dengue virus is responsible for approximately 100-400 million infections worldwide each year (1,2), with severe cases leading to dengue hemorrhagic fever and dengue shock syndrome. Zika virus, on the other hand, gained global attention in 2015 due to rapid expansion across the tropical and subtropical countries in the Americas and more importantly its association with neurological symptoms and congenital malformations in newborns (3,4). Both viruses are primarily transmitted through the bite of infected *Aedes* mosquitoes. At present, most research on DENV and ZIKV transmission gives primary focus to *Ae. aegypti* as the main vector and *Ae. albopictus* as a secondary one, while other mosquito species are largely overlooked. However, with climate change, there’s potential for arbovirus vectors to spread geographically (5–7), including neglected mosquito species posing an unknown risk of virus transmission. Additionally, several species-specific intervention such as gene drive and *Wolbachia* might cause significant changes in population structure of the primary vectors thus providing ecological gap for the neglected vectors.

The Scutellaris group of *Aedes* consists of more than 46 species (8) with geographical range originally covers the Southeast Asia, South Pacific, and Northern Australia (9–11). The major species of this subgroup is *Ae. albopictus*, which has been one of the most invasive mosquitoes globally. *Ae. scutellari*s (Walker, 1859), also belong to this subgroup, has been considered a potential carrier of the dengue virus in Papua New Guinea. Early reports suggested *Ae. scutellaris* to be responsible for major DENV transmission in the Pacific islands (10,11). *Ae. scutellari*s geographic range covers Papua New Guinea, Tonga, Southeast Asia, the South Pacific, Australia, and central Thailand (10,12,13). Despite the suggested role of *Ae. scutellaris* in DENV transmission, only one study has made a comparative analysis of vector competence regarding DENV2 infection in *Ae. scutellaris* and the primary vector *Ae. Aegypti* (10). The study found that *Ae. scutellaris* is a moderately efficient DENV2 vector, with salivary gland infection to those of *Ae. aegypti*. However, the study only involved DENV2 without evaluation of the virus’s ability to escape from the salivary gland into the mosquito’s saliva as the infective stage.

In this present study, we conducted detailed analyses of DENV1-4 and ZIKV infection kinetics to determine the vector competence of *Ae. scutellaris*, and compared it to Thai laboratory colonies of *Ae. aegypti* and *Ae. albopictus*. Through a combination of artificial membrane feeding and intrathoracic injection, we demonstrated that *Ae. scutellaris* can transmit ZIKV and DENVs at a level similar to *Ae. aegypti*, especially inoculated with the same virus load.

## 2. Materials and Methods

### 2.1 Ethic statement

This study was carried out in accordance with the Faculty of Tropical Medicine-animal Care and Use Committee (FTM-ACUC), Mahidol University, Bangkok and the BIOTEC Committee for use and care of laboratory animals. Mosquito collection and maintain the field colony was processed followed by the approved protocol of FTM – ACUC 008/2023 and BT-Animal 05/2564. Mice were used only for mosquito rearing as a blood source, according to the approved protocol (BT-Animal 05/2564). Mosquito infection assays were performed followed the approved protocol (BT-Animal 05/2564). Human erythrocytes were used for an infection by artificial membrane feeding. Blood was collected from human volunteer following the approved protocol (NIRB-052-2563).

### 2.2 *Aedes scutellaris* colonization

*Aedes scutellaris* subgroup larvae were collected from the coastal area at Song Khlong Subdistrict, Bang Pakong District, Chachoengsao Province between November 2022 to January 2023 (**Figure 1**). A total of approximately 1600 *Aedes spp.* larvae were collected from the field sites. These larvae were subsequently transferred to an insectarium at Faculty of Tropical Medicine, Mahidol university and reared to adults. Individual emerging of *Ae. scutellaris* identified by morphological observation (14) were pooled to establish a field-derived *Ae. scutellaris* colony. Molecular identification of *Ae. scutellaris* was confirmed by PCR based DNA barcode of COI as previously described (Supplementary document 1) (15). Low passage *Ae. scutellaris* (less than six generations) were transferred to BIOTEC’s insectary and used for infection experiments.

**Figure 1.**
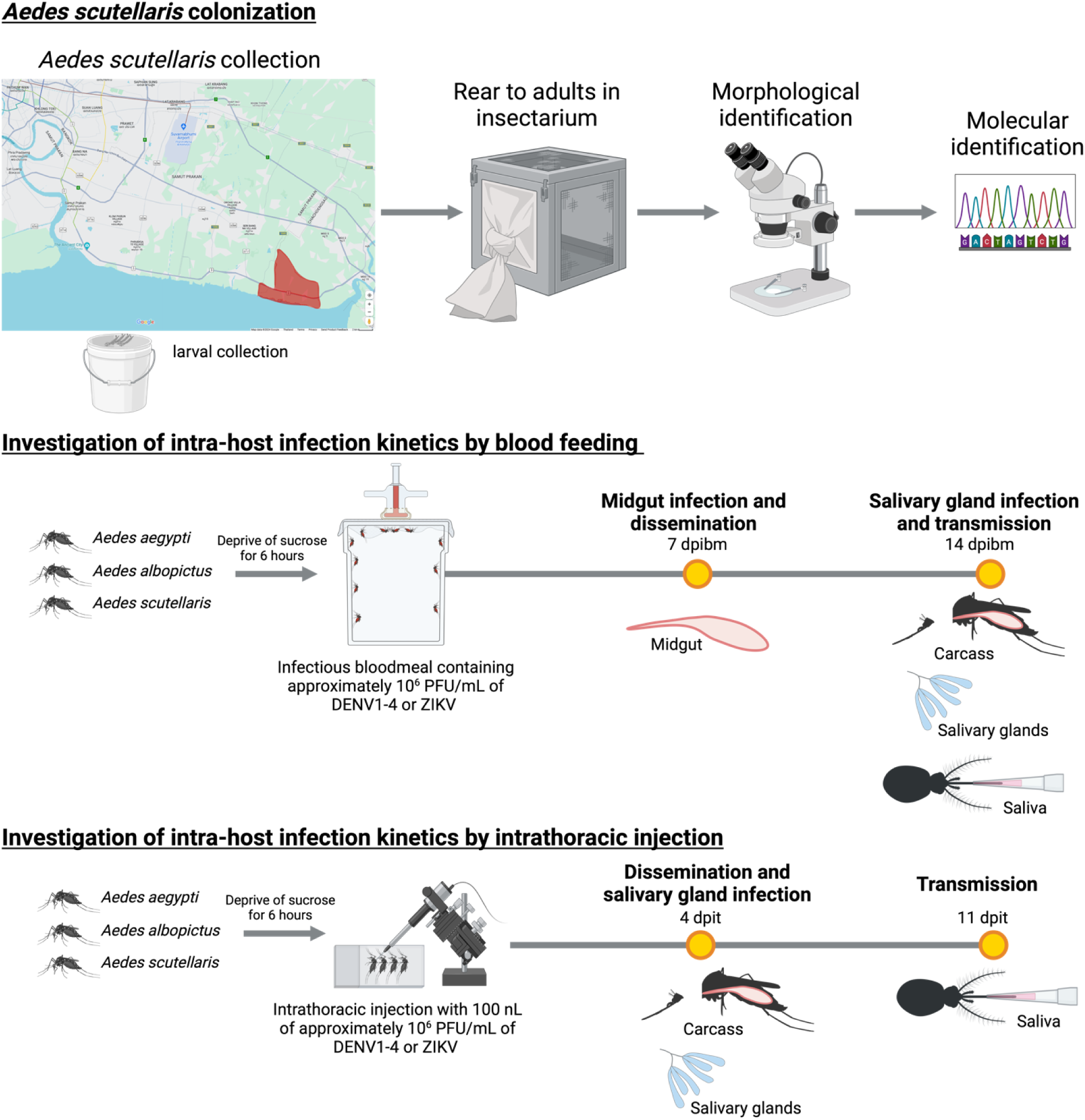
Overview of the *Aedes scutellaris* colonization and arbovirus infection kinetics investigation.

### 2.3 Animal maintenance

The recently colonized *Ae. scutellaris* and laboratory strains of *Ae. aegypti* and *Ae. albopictus* obtained from Department of Medical Sciences, Ministry of Public Health, Thailand were used for all the infection studies. Mosquitoes were maintained in BIOTEC’s insectary at 27 °C with 80% humidity and a 12-hours day/night, 30-minutes dusk/dawn lighting cycle. The larvae were fed on powdered fish food (Tetra Bits). Adults were fed on 10% sucrose solution ad libitum. To obtain the eggs for colony maintenance, mosquitoes were allowed to feed on ICR mice anesthetized with 2% Avertin (2,2,2-Tribromoethanol, Sigma, T48402).

### 2.4 Virus propagation and titration

The SV0010/15 Thai ZIKV isolate and the contemporary DENV 1-4 panel from BEI Resources were used in the infection studies. All the viruses were propagated using the *Ae. albopictus* cell line C6/36 (ATCC CRL-1660) as previously described (16). Briefly, after the cells were cultured to 80% confluency in T-75 cm^2^ flasks,supernatant was removed and replaced with the virus stocks at the MOI of 0.1 in 5 mL incomplete L-15 medium for 2 hours. After virus incubation, supernatant was removed, and replaced with 2% FBS L-15 medium then further incubated at 28 °C. The supernatant was collected at 6-7 days post inoculation then supplemented with FBS to final concentration of 20% and stored at 80 °C until further use.

Virus titers was determined by plaque assay in BHK-21 cells following the previously published protocol (16). Briefly, 100 µL of virus suspension was added to BHK-21 cells seeded in 24-well plate at 80% confluency. Inoculated plates were then gently rocked at room temperature for 15 minutes before an incubation at 37 °C, 5% CO_2_ for 45 minutes. After incubation, 1 mL of overlay medium (1% methylcellulose (Sigma, M0512) in MEM supplemented with 2% FBS and 1X Pen/Strep) was added to each well then further incubated at 37 °C, 5% CO_2_ for 6 days. The plates were then fixed and stained with 0.5% crystal violet (Sigma, C6158) in 1:1 Methanol/Acetone fixative for 1 hour at room temperature. Stained plates were then washed under running tap water and air dry before plaque counting.

### 2.5 Mosquito infection by artificial membrane feeding

Mosquitoes were orally challenged with ZIKV and DENVs using the Hemotek artificial membrane feeding system. Briefly, 7-day-old female mosquitoes were deprived of sucrose solution overnight before offering with an artificial infectious blood meal containing 40% washed human erythrocytes and virus stock diluted to approximately 6 Log10 PFU/mL for 30 minutes. The measured exact feeding titers for each virus were 6.26-6.54 log_10_ PFU/mL for DENV1, 6.32-6.38 log_10_ PFU/mL for DENV2, 6.40-6.41 log_10_ PFU/mL for DENV3, 6.08-6.34 log_10_ PFU/mL for DENV4, and 6.32-6.56 log_10_ PFU/mL for ZIKV. After feeding, mosquitoes were then anesthetized in a refrigerator for 15 minutes, and engorged females were sorted on ice. Blood-fed mosquitoes were maintained in waxed paper cups with 10% sucrose solution in a climate-controlled chamber under controlled conditions of 28 °C ± 1 °C, a 12-hour light-dark cycle, and 70% relative humidity before tissue collection.

### 2.6 Estimation of bloodmeal size

Blood meal size of the blood engorged mosquitoes was estimated by measuring the amount of heme. The recently blood-engorged mosquitoes (approximately 30 minutes after offering the blood meal) were collected in 100 µL sterile Milli-Q water and stored at -80 °C until use. Blood meal used for blood feeding was also stored at -80 °C to use as a standard. To measure heme amount, the samples were thawed and homogenized with 0.5 mm glass beads using Bullet Blender Tissue Homogenizer (NextAdvance). Supernatant of the homogenized mosquitoes was then collected after centrifugation at 8,000 xg, 4 °C for 2 minutes. Amount of heme in the supernatant was then measured using Heme assay kit (Sigma MAK316) following the manufacturer’s protocol. Briefly, 50 µL of supernatant was added to 96-well plate containing 50 µL of Milli-Q water and 200 µL of heme assay reagent, incubate at room temperature for 5 minutes then measure the absorbance at 400 nm. For the standard, 6 adult female mosquitoes were homogenized in 600 µL Milli-Q water and 50 µL of supernatant was added to 96-well plate containing 0, 1, 2, 3, 4, or 5 µL of blood meal, Milli-Q water up to 50 µL, and 200 µL of heme assay reagent. Estimated blood meal size was then calculated using the linear standard curve.

### 2.7 Mosquito infection by intrathoracic injection

The intrathoracic inoculation was conducted by injected 100 nL of virus stock into the thorax of cold-anesthetized 4-to 7-day-old female mosquitoes using a nanoliter injector (Nanoject III; Drummond Scientific). The measured titers used for injection for each virus were 6.23-6.40 log_10_ PFU/mL for DENV1, 6.11-6.34 log_10_ PFU/mL for DENV2, 5.85-6.40 log_10_ PFU/mL for DENV3, 6.11-6.41 log_10_ PFU/mL for DENV4, and 6.36-6.48 log_10_ PFU/mL for ZIKV. Injected mosquitoes were then maintained on 10% sucrose solution at a condition as mentioned above.

### 2.8 Mosquito dissection and salivation assay

Mosquitoes were cold anesthetized in refrigerator for 15 minutes before surface sterilization in 70% ethanol for 1 minute followed by twice PBS washes. Mosquitoes were then individually dissected in drops of 1X PBS. Midguts, carcasses, and salivary glands were collected in 150 µL of MEM medium supplemented with 10% FBS and 1X Pen/Strep and stored at -80 °C for titration with plaque assay as described above.

Mosquito saliva was collected according to a previously published protocol (16). Briefly, mosquitoes were paralyzed with triethylamine before inserting the proboscis into a pipette tip containing 20 µL of MEM supplemented with 10% FBS and 1X Pen/Strep. After 45 minutes of salivation, the medium in the tips were mixed with 180 µL of MEM supplemented with 2% FBS and 1X Antibiotics/Antimycotics and immediately titrated by plaque assay mosquito.

### 2.9 Data analysis

The Factor Analysis of Mixed Data (FAMD) (17) was used to identify relationship between infection prevalence/median from each infection experiment and mosquito species, virus, and tissue type. FAMD was conducted using FactoMineR package in R (18).

Statistical analyses in this study were conducted using the rstatix package (version 0.7.1) (19) in R (version 4.3.0). Multiple comparison was conducted using Kruskal-Wallis followed by Dunn’s posthoc test. Graphs were generated using the ggpubr package (version 0.6.0) (20) in R.

## 3. Results

In order to determine the vector competence of *Ae. scutellaris* for dengue virus serotypes 1-4 (DENV1-4) and Zika virus (ZIKV), we conducted a study to evaluate the intra-host infection kinetics of these viruses in a recently colonized *Ae. scutellaris* population. We compared the infection levels of *Ae. scutellaris* with those of laboratory colonies of *Ae. aegypti* and *Ae. albopictus* with high arbo-flavivirus transmissibility. The mosquitoes were fed a blood meal containing approximately 6 log10 PFU/mL of each virus. We assessed the extent of midgut infection 7 days post the infectious blood meal (dpibm), and then determined the dissemination, salivary gland infection, and transmissibility 14 dpibm (**Figure 1**). Additionally, due to a low engorgement rate of *Ae. scutellaris* from artificial membrane feeding, we also infected the mosquitoes by intrathoracic inoculation to determine vector competence when the mosquitoes were infected with the same amount of inoculating viruses. Virus replication in the body and salivary gland infection were determined at 4 days post intrathoracic injection (dpit), and transmissibility was determined at 11 dpit (**Figure 1**).

### 3.1 *Aedes scutellaris* exhibits a lower prevalence of midgut infection following artificial membrane feeding, but had similar level of transmissibility compared to *Ae. aegypti*

The arbo-flavivirus susceptibility and transmissibility of each *Aedes* species was compared as demonstrated by infection prevalence through each infection barrier (percent of mosquito with infectious virus specific body compartment in total blood fed mosquitoes). The Factor analysis of mixed data (FAMD) was used to investigate the relationship of infection prevalence pattern between different factors mosquito species, tissues, and virus. We found that the prevalence of infection of *Ae. scutellaris* was similar to *Ae. albopictus*, both of which differ from *Ae. aegypti* (**Figure 2A**). To compare overall arbo-flaviviruses susceptibility of each mosquito, we next compared infection prevalence by grouping the data according to mosquito species (**Figure 2B-E**, left panel).

**Figure 2.**
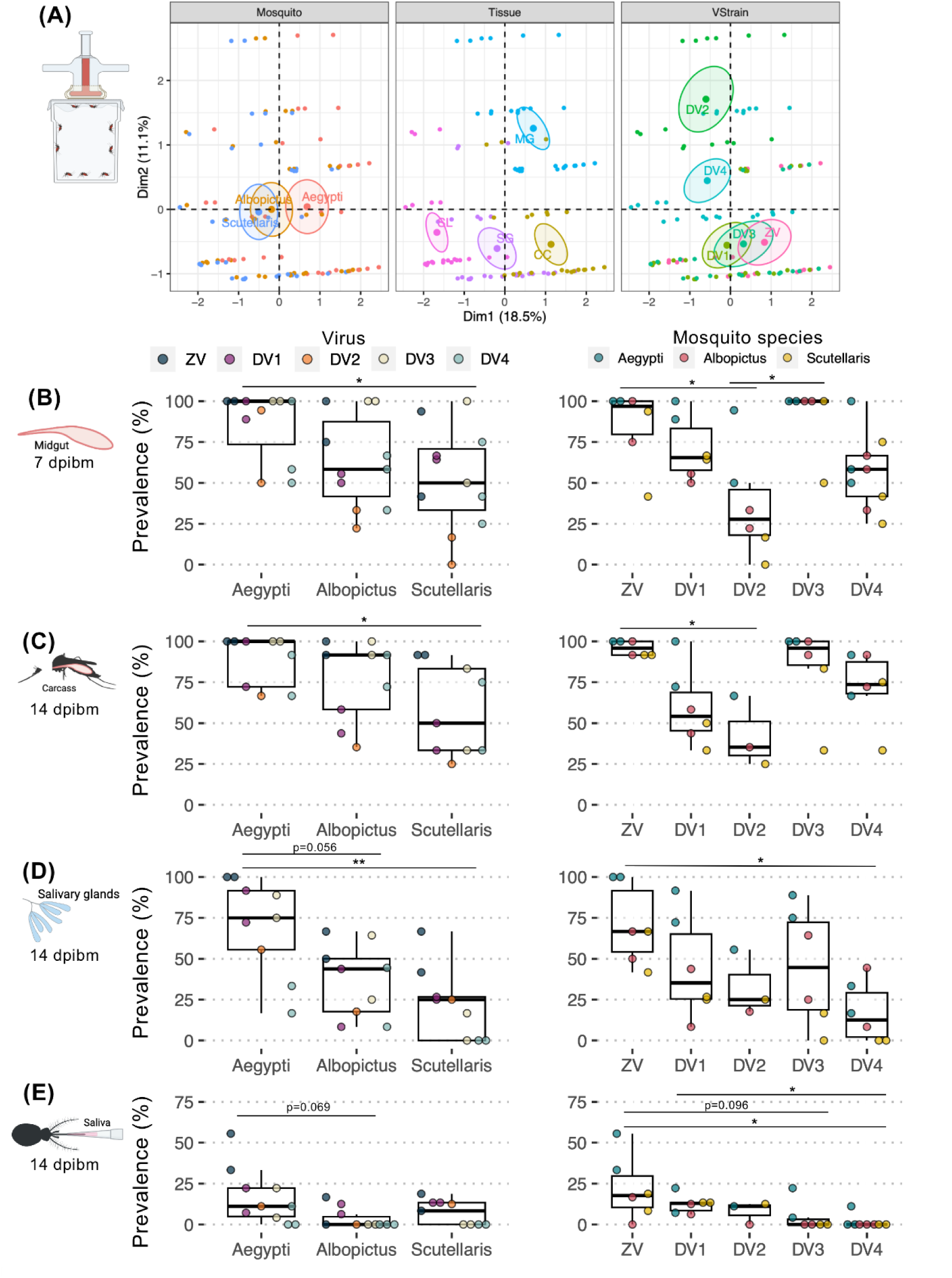
Comparison of ZIKV and DENV1-4 infection prevalence in different infection barriers of *Ae. aegypti*, *Ae. albopictus*, and *Ae. scutellaris* after artificial membrane feeding. (A) Factor Analysis of Mixed Data to visualize relationship between infection prevalence results from each blood feeding experiment on the transformed coordinates. The information in the analysis included infection prevalence, mosquito species (Mosquito), tissues type (Tissue), and virus (VStrain). The clustering is based on qualitative coordinates (mosquito species, tissue and virus), and each color represents individuals in a specific feature. Cluster distance represents the correlation among variables. The confidence ellipses indicate their potential relationships. Abbreviation: Dim, dimension. Box and scatter plot comparing infection prevalence in the (B) midgut at 7 dpibm, (C) carcass at 14 dpibm, (D) salivary glands at 14 dpibm, and (E) saliva at 14 dpibm grouping by mosquito species or virus. Each dot represents infection prevalence of from each blood feeding experiment. *: p<0.05, **: p<0.01.

The midgut overall arbo-flaviviruses (ZIKV and DENV1-4) infection prevalence were the lowest in *Ae. scutellaris* followed by *Ae. albopictus,* and lastly *Ae. aegypti* with the median midgut infection prevalence of 50, 58.3, and 100 %, respectively (**Figure 2B**, left panel). Given that the size of blood meal directly influences the inoculation size, it was possible that the observed differences in establishment of midgut infection were due to the differences in blood meal size. Therefore, we determined blood meal size by quantifying amount of heme in the blood engorged mosquitoes. We found that *Ae. aegypti*. *Ae. albopictus*, and *Ae. scutellaris* ingested 3.0 ± 0.4, 2.5 ± 0.7, and 1.6 ± 0.6 µL of blood, respectively (**Figure 3**). With the largest blood meal size among the three *Aedes* species, it was not surprising that *Ae. aegypti* had the highest midgut infection prevalence at 7 dpibm. Interestingly, despite *Ae. scutellaris* having a significantly smaller blood meal size than *Ae. albopictus*, the midgut infection prevalence was similar, suggesting a more permissive midguts of *Ae. scutellaris* compared to *Ae. albopictus*.

**Figure 3.**
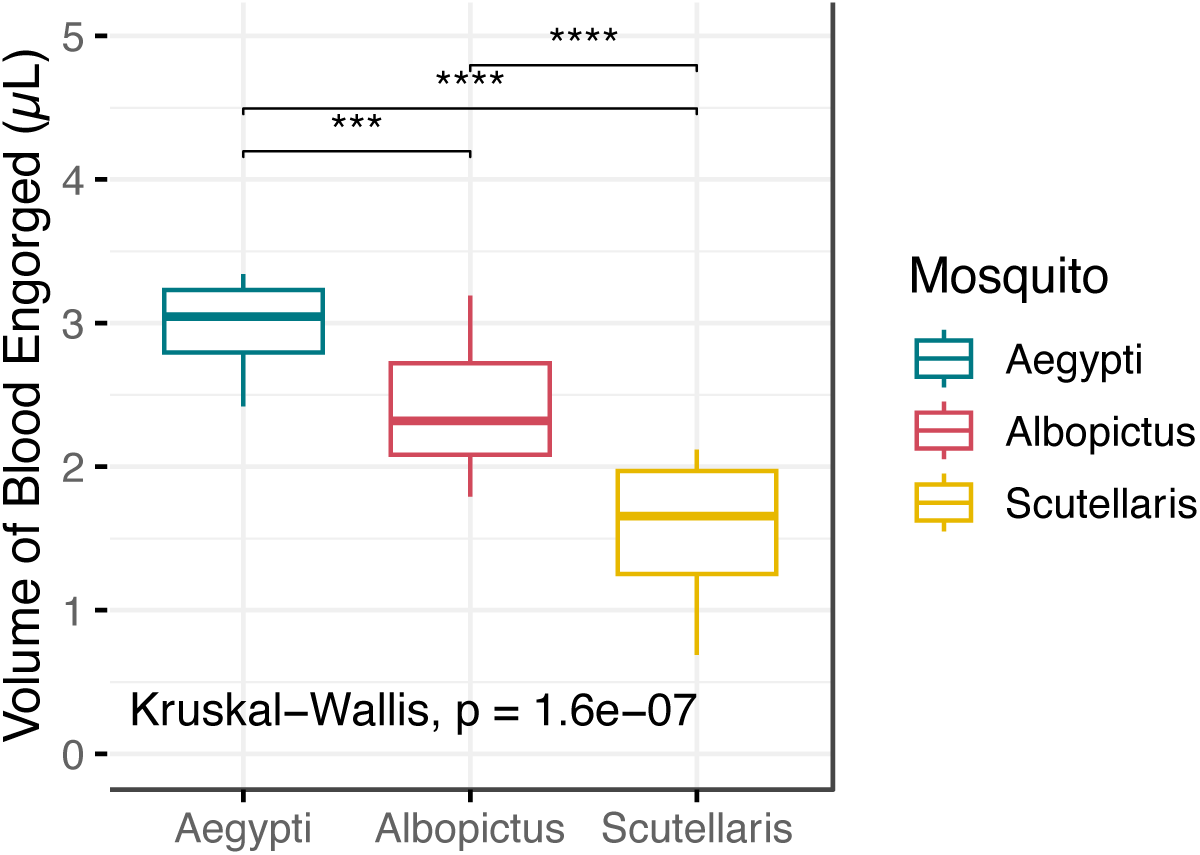
Volume of bloodmeal engorged by *Ae. aegypti*, *Ae. albopictus*, and *Ae. scutellaris*. Box and scatter plot summarizing bloodmeal volume engorged by individual mosquito. Each dot represents bloodmeal volume engorge by each mosquito. ***: p<0.001, ****: p<0.0001.

Subsequently in the transmission cycle, the carcasses and salivary glands infection prevalence among three *Aedes* species followed the same pattern as those in the midguts, with *Ae. aegypti* being the most permissible while *Ae. scutellaris* being the least permissible mosquitoes (**Figure 2C-D**, left panel). In salivary glands, the median infection prevalence of all arbo-flaviviruses in *Ae. aegypti*. *Ae. albopictus*, and *Ae. scutellaris* were 75%, 50% and 25%, respectively. This demonstrates that the establishment of midgut infection is an important bottleneck that dictate the successful infection in subsequent steps of transmission cycle. Interestingly, despite lower salivary gland prevalence in *Ae. scutellaris* compared to *Ae. aegypti*, the transmissibility of arbo-flaviviruses between these two species were not statistically significant with the median prevalence of 11.1 % for *Ae. aegypti* and 8.3% for *Ae. scutellaris* (**Figure 2E**, left panel). On the contrary, the median prevalence in saliva of *Ae. albopictus* was 0%. These results demonstrate minimal salivary gland escape barrier of *Ae. scutellaris* but strong salivary gland escape barrier of *Ae. albopictus*.

### 3.2 Tissue-specific infectivity of each arbo-flavivirus was largely determined by viral genetics

In addition to visualizing relationship of infection prevalence data according to mosquito, the FAMD clustering was also conducted based on virus strain. We found that the pattern of infection prevalence phenotype of ZIKV, DENV1 and DENV3 were clustered together while those of DENV2 and DENV4 had distinct clusters (**Figure 2A**).

To compare overall infectivity of each virus in detail, the infection prevalence was analyzed grouping by virus (**Figure 2B-E**, right panel). Among all the arbo-flaviviruses tested, ZIKV and DENV3 displayed the highest midgut infection prevalence across all three *Aedes* species with the median midgut infection prevalence of 96.9 and 100%, respectively (**Figure 2B**, right panel). The other DENVs exhibited varying levels of midgut infection prevalence with the median prevalence across all mosquito species of 65.5%, 27.8%, and 58.3%, for DENV1, DENV2, and DENV4 respectively. These results demonstrate that ZIKV and DENV3 were efficient at establishing primary midgut infection while DENV2 had the poorest establishment of midgut infection.

When comparing between viruses, our results demonstrate superior body and salivary gland infectivity as well as transmissibility of ZIKV over DENVs across all three *Aedes* species followed by DENV1 (**Figure 2B-E**, right panel). Interestingly, while DENV2 had the lowest midgut infection prevalence, the virus has minimal barrier during subsequent infection steps resulting in similar transmissibility to DENV1 (**Figure 2B-E**, right panel) demonstrating low intra-host infection barrier of DENV2 when crossing barrier. On the contrary, while DENV3 exhibited the highest midgut infection prevalence, those of the subsequent steps decreased to a level similar to other DENVs suggesting that DENV3 infection was less efficient during subsequent intra-host barrier (**Figure 2B-D**, right panel). In fact, DENV3 was among the viruses with lowest transmissibility (**Figure 2E**, right panel). In addition to DENV3, DENV4 also exhibited strong barrier during both salivary gland infection and escape barriers (**Figure 2D-E**, right panel). The tissue-specific infectivity phenotypes of ZIKV and DENVs were conserved among the three *Aedes* species demonstrating that the infectivity was largely determined by virus genetics.

### 3.3 Tissues of all three *Aedes* species support high level of arbo-flavivirus replication

In addition to the infection prevalence, we also investigated how well the tissues of *Ae. scutellaris* could support arbo-flaviviruses replication **Figure 4**. The risk of arbo-flaviviruses transmission is likely to increase with higher replication rates in the insect vectors. In this analysis, we compared the levels of infectious virus in each body compartment of infected mosquitoes. The FAMD map suggested that the median virus titers in the infected *Ae. aegypti* was more similar to *Ae. albopictus* than *Ae. scutellaris*, which was different from the FAMD map of the infection prevalence that demonstrate more similarity between *Ae. albopictus* and *Ae. scutellaris*. The differences between the FAMD map of infection prevalence and titers was also observed among the virus cluster. While the infection prevalence of DENV1, DENV3 and ZIKV were clustered together, only ZIKV and DENV1 were clustered together and the DENV3 became more similar to DENV2 and DENV4 (**Figure 4A**, **Supplementary Figure S1**).

**Figure 4.**
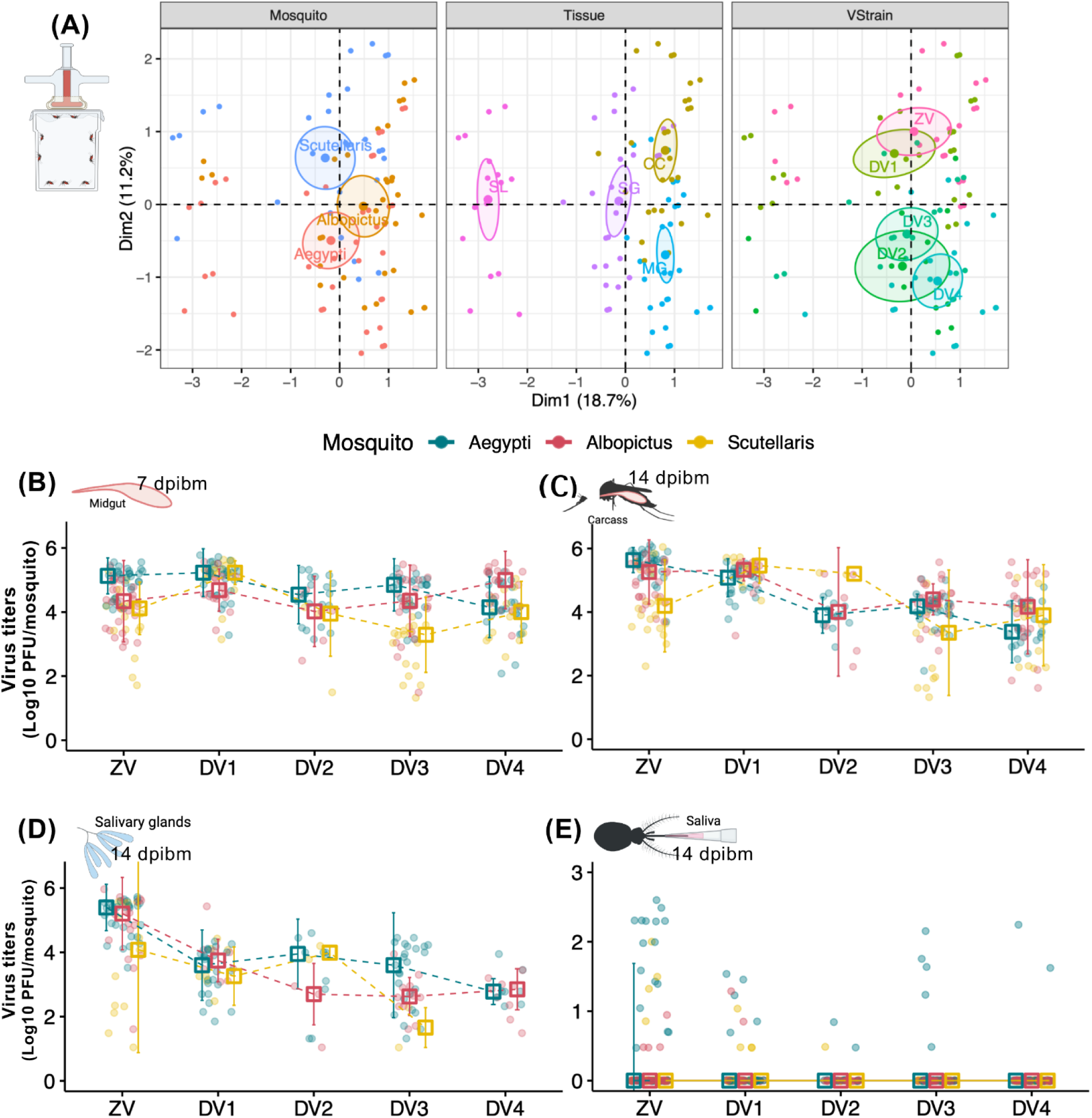
Comparison of ZIKV and DENV1-4 titers in different infection barriers of *Ae. aegypti*, *Ae. albopictus*, and *Ae. scutellaris*. (A) FAMD factor maps to visualize variance between median infection titers from each blood feeding experiment on the transformed coordinates. The information in the analysis included median infection titers, mosquito species (Mosquito), tissues type (Tissue), and virus (VStrain). The clustering is based on qualitative coordinates (mosquito species, tissue and virus), and each color represents individuals in a specific feature. Cluster distance represents the correlation among variables. The confidence ellipses indicate their potential relationships. Abbreviation: Dim, dimension. Box and scatter plot comparing infection titers of individual mosquito in the (B) midguts at 7 dpibm, (C) carcasses at 14 dpibm, (D) salivary glands at 14 dpibm, and (E) saliva at 14 dpibm grouping by mosquito species and virus. Each dot represents infection prevalence of from each blood feeding experiment. *: p<0.05, **: p<0.01.

Further comparison of the midgut infection revealed that titers of ZIKV and DENV3 in infected *Ae. scutellaris* were significantly lower than in *Ae. aegypti*, but not for DENV1, DENV2, and DENV4 (**Figure 4B**, **Supplementary Figure S1B**). These findings indicate that *Ae. scutellaris* midguts can support similar levels of DENV1, DENV2, and DENV4 during the late stage of midgut infection, despite a smaller starting inoculum from a smaller blood meal size. Similarly, midgut titers of ZIKV in *Ae. albopictus* was significantly lower than *Ae. aegypti* while those of DENVs were similar among these two *Aedes* species or even higher than *Ae. aegypti* in the case of DENV4 (**Figure 4B**, **Supplementary Figure S1B**).

In the next infection barrier, the pattern of infection intensity (virus titers) in the carcasses was similar to those of the midgut infection levels, with *Ae. scutellaris* displaying comparable DENV2 and DENV4 titers, as well as lower ZIKV and DENV3 titers compared to *Ae. aegypti* (**Figure 4C**, **Supplementary Figure S1C**). Interestingly, DENV1 titers in the carcasses of *Ae. scutellaris* were significantly higher than those in *Ae. aegypti*, suggesting a higher level of DENV1 replication in *Ae. scutellaris* during the late stage of infection. In *Ae. albopictus*, the carcass titers of DENV1-4 were not different from *Ae. aegypti* but those of ZIKV were slightly lower.

The titers of each virus in the infected salivary glands were not statistically different among the three *Aedes* species (**Figure 4D**, **Supplementary Figure S1D**), indicating that these viruses can efficiently replicate in the salivary glands of *Ae. scutellaris* once infection is established. However, it should be noted that lack of significance can also possibly be due to the low number of infected salivary glands. Indeed, none of the *Ae. scutellaris* salivary glands were infected by DENV4.

### 3.4 *Aedes aegypti* exhibits superior arbo-flavivirus transmissibility than *Ae. albopictus* and *Ae. scutellaris* after artificial membrane feeding

The virus titers of ZIKV and DENV3 in *Ae. aegypti* saliva was significantly higher than in the other two *Aedes* species **Figure 4E**. For DENV1, DENV2 and DENV4, although the virus titers in *Ae. aegypti* saliva was higher than the other two *Aedes* species, the number of mosquitoes with the virus in their saliva was low making it difficult to draw conclusion regarding their transmissibility.

### 3.5 *Aedes scutellaris* is as robust as *Aedes aegypti* in arbo-flavivirus transmission when inoculated with similar virus titers

Since *Ae. scutellaris* took a significantly smaller blood meal from the artificial membrane feeding, it is possible that the smaller starting inoculating virus affect the differential transmissibility phenotype among three *Aedes* species. Such effect from different amount of inoculating virus on transmissibility has been demonstrated, even with the same virus and mosquito strain in our previous detailed ZIKV infection kinetics study (16). It is possible that the recently colonized *Ae. scutellaris* is highly anthropophilic, thus the blood meal size from artificial membrane feeding was underestimated, which mosquitoes in the field might take larger blood meal than what we observed in the laboratory resulting in an underestimation of *Ae. scutellaris* transmissibility. To overcome this limitation, we infected ZIKV or DENVs in the three *Aedes* species by intrathoracic injection to ensure equal virus inoculation. Here, each individual mosquitoes were injected with approximately 1,000 PFU/mL of virus. The body and salivary gland titers were measured at 4 dpit and transmission was evaluated by collecting saliva at 11 dpit.

The FAMD analysis revealed that the clusters of *Ae. scutellaris* and *Ae. aegypti* were almost identical suggesting similar infection prevalence patterns between the two mosquitoes (**Figure 5A**). In contrast to the blood feeding experiment that demonstrated a lower infection and transmissibility of *Ae. scutellaris* compared to *Ae. aegypti*, we found that the infection prevalence of arbo-flaviviruses of these two *Aedes* species were comparable throughout the mosquito transmission cycle (**Figure 5B-D**, left panel). While *Ae. albopictus* had similar infection prevalence in the body and salivary glands to the other two *Aedes* species, the prevalence of arbo-flaviviruses in the saliva of *Ae. albopictus* was 21.4% compared to 53.3% and 60% in *Ae. aegypti* and *Ae. scutellaris*, respectively (**Figure 5D**, left panel). This demonstrated a stronger salivary gland escape barrier in *Ae. albopictus*.

**Figure 5.**
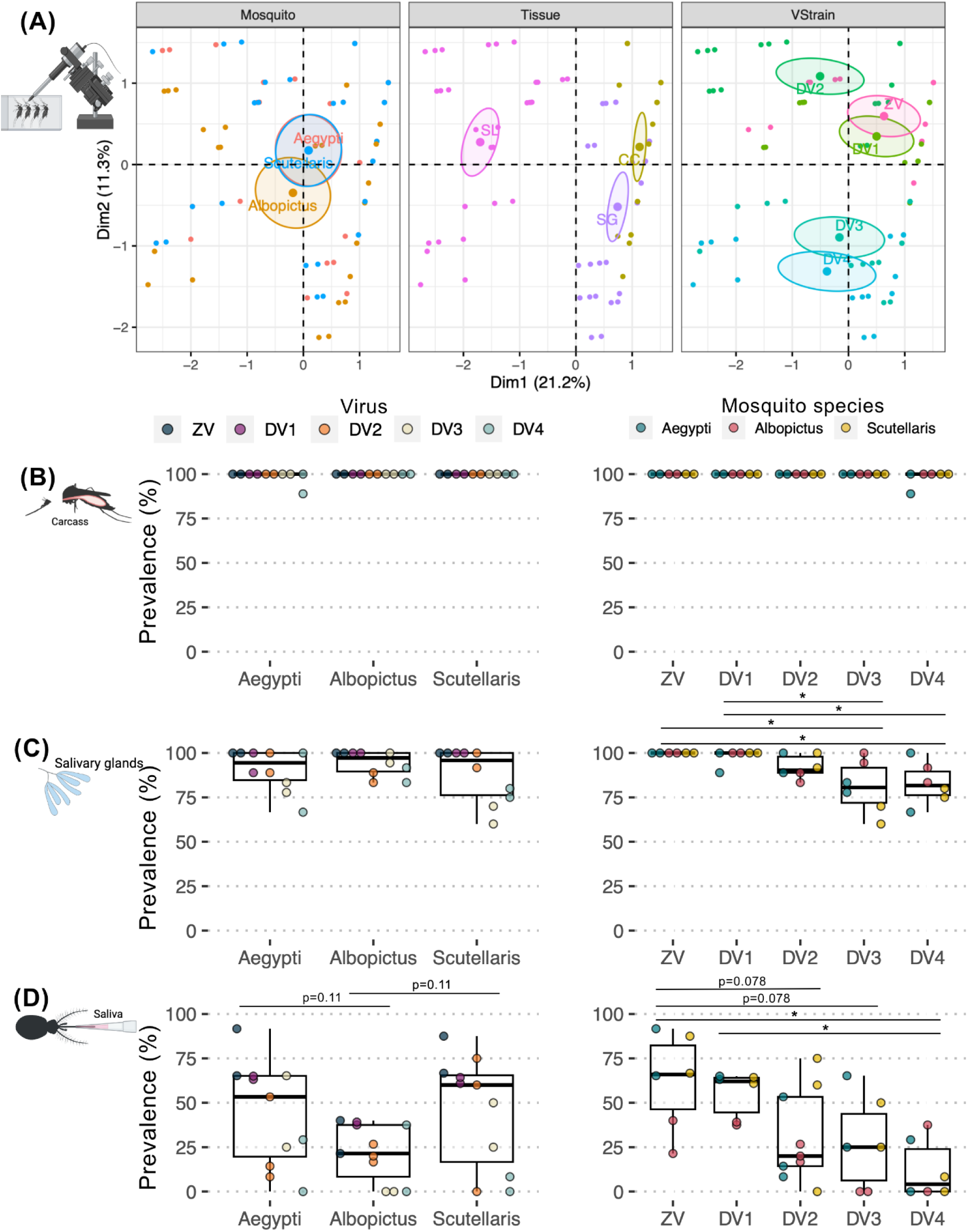
Comparison of ZIKV and DENV1-4 infection prevalence in different infection barriers of *Ae. aegypti*, *Ae. albopictus*, and *Ae. scutellaris* after intrathoracic injection. (A) FAMD factor maps to visualize variance between median infection titers from each blood feeding experiment on the transformed coordinates. The information in the analysis included median infection titers, mosquito species (Mosquito), tissues type (Tissue), and virus (VStrain). The clustering is based on qualitative coordinates (mosquito species, tissue and virus), and each color represents individuals in a specific feature. Cluster distance represents the correlation among variables. The confidence ellipses indicate their potential relationships. Abbreviation: Dim, dimension. Box and scatter plot comparing infection titers of individual mosquito in the (B) carcasses at 4 dpit, (C) salivary glands at 4 dpit, and (D) saliva at 11 dpit, grouping by mosquito species and virus. Each dot represents infection prevalence of from each intrathoracic injection experiment. Statistical analysis comparing infection level was conducted using Kruskal-Wallis followed by Dunn’s posthoc test in R. *: p<0.05, **: p<0.01.

Unlike the infection prevalence, the FAMD map of the median titers demonstrated that the pattern of infection titers of the three *Aedes* species were different (**Figure 6A**). When comparing the infection titers to *Ae. aegypti*, we found that *Ae. scutellaris* supported lower levels of ZIKV replication in the body and salivary glands but exhibited similar ZIKV tiers in the saliva (**Figure 6B-D**) suggesting minimal ZIKV salivary gland escape barrier of *Ae. scutellaris*. Additionally, we found that all DENV propagated to a similar or higher level in *Ae. scutellaris* than in *Ae. aegypti* after intrathoracic inoculation and eventually resulted in similar saliva virus titers (**Figure 6B-D**). While *Ae. albopictus* had the most robust virus propagation in the body and especially the salivary glands (**Figure 6A-B**), the mosquito had the lowest DENVs titers in the saliva compared to the other two *Aedes* species (**Figure 6C**).

**Figure 6.**
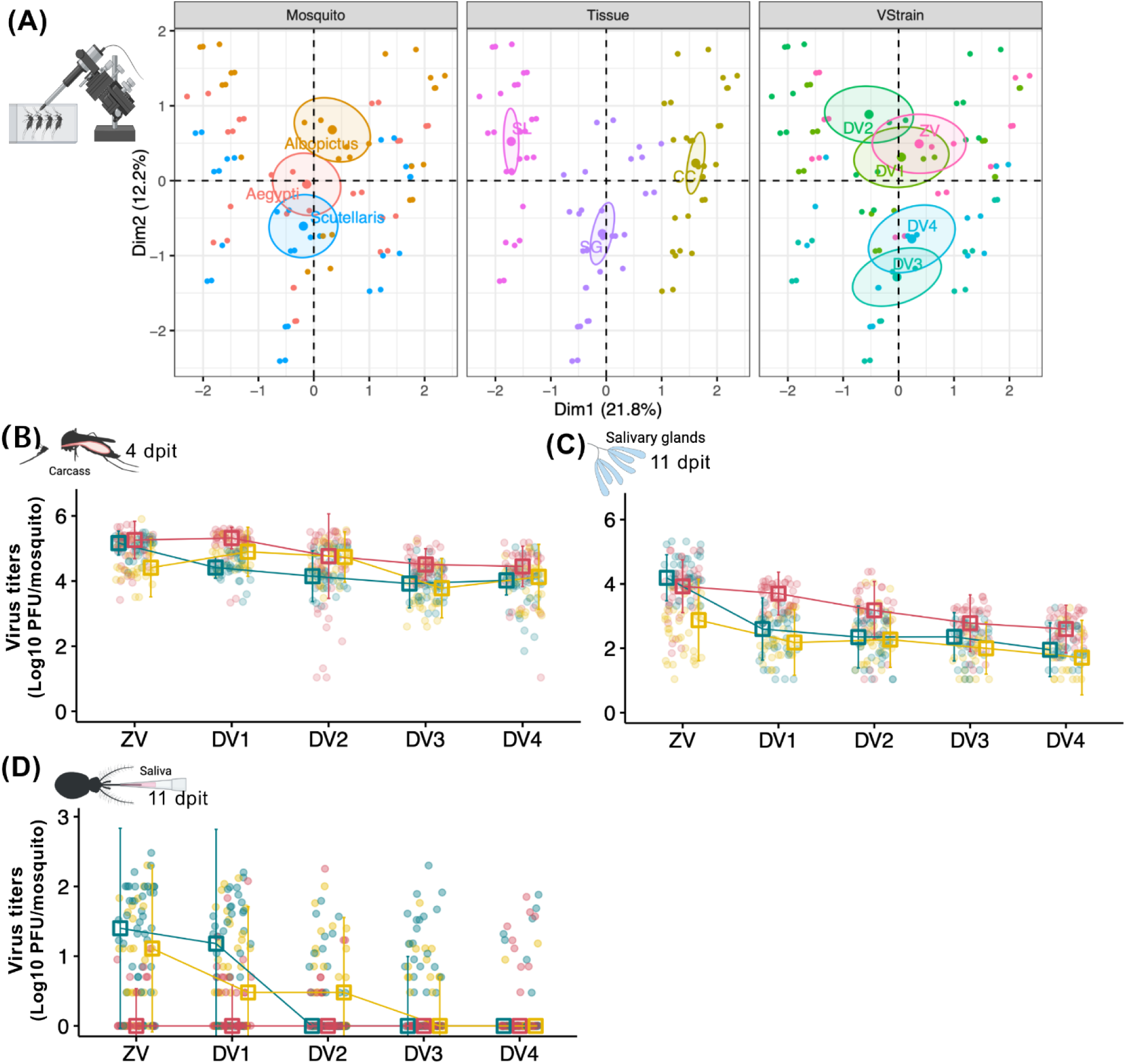
Comparison of ZIKV and DENV1-4 titers in different infection barriers of *Ae. aegypti*, *Ae. albopictus*, and *Ae. scutellaris* after intrathoracic injection. (A) FAMD factor maps to visualize variance between median infection titers from intrathoracic injection experiment on the transformed coordinates. The information in the analysis included median infection titers, mosquito species (Mosquito), tissues type (Tissue), and virus (VStrain). The clustering is based on qualitative coordinates (mosquito species, tissue and virus), and each color represents individuals in a specific feature. Cluster distance represents the correlation among variables. The confidence ellipses indicate their potential relationships. Abbreviation: Dim, dimension. Box and scatter plot comparing infection titers of individual mosquito in the (B) carcasses at 4 dpit, (C) salivary glands at 4 dpit, and (D) saliva at 11 dpit, grouping by mosquito species and virus. Each dot represents infection prevalence of from each intrathoracic injection experiment. *: p<0.05, **: p<0.01.

### 3.5 ZIKV superior infectivity and transmissibility compared to DENVs were observed across all three *Aedes* species

Although **Figure 2** shows ZIKV’s higher infectivity after passing midgut infection compared to DENVs, it is possible that the strong midgut infection barriers lead to the lower infection level during subsequent transmission cycle. The intrathoracic injection results confirmed that, even when bypassing the midgut infection, ZIKV were among the highest infection prevalence and titers compared to DENVs, regardless of the tissue and mosquito species (**Figure 5** and **Figure 6**).

## 4. Discussion

To evaluate the potential role of *Ae. scutellaris* in DENV and ZIKV transmission, we compared the intra-host infection kinetics of these viruses in a recently colonized *Ae. scutellaris* with those of laboratory colonies of *Ae. aegypti* and *Ae. albopictus*, which are known to have high arbo-flavivirus transmissibility. In fact, the laboratory *Ae. aegypti* strain was the most transmissible mosquito we have in our laboratory. The comparison to transmissibility of *Ae. scutellaris* to this highly transmissible strain thus highlight the transmission potential of *Ae. scutellaris*. Our research provides extensive insights into the vector competence of *Ae. scutellaris* for dengue and Zika viruses in relation to the more studied *Ae. aegypti* and *Ae. albopictus*, offering a deeper understanding of the potential role this mosquito species plays in the spread of these diseases.

Our results demonstrate that the size of blood meal (which also determine the size of initial inoculum) plays an important role in determining the establishment of midgut infection, a critical barrier for arbovirus transmission. With the largest size of ingested blood, *Ae. aegypti* had the highest midgut infection prevalence among the three *Aedes* species. Despite *Ae. scutellaris* having a significantly smaller blood meal size than *Ae. albopictus*, the midgut infection prevalence was similar, suggesting a more permissive midguts of *Ae. scutellaris* compared to *Ae. albopictus*. Although the current work did not identify which step of midgut infection posed significant bottleneck for *Ae. scutellaris* and *Ae. albopictus*, the fact that the midgut virus titers eventually reached similar levels to the other two *Aedes* species at the late timepoint suggested that the barrier occur s during primary midgut infection.

Although not surprising, our study provided evidence on the effect of bloodmeal size on vector competence. The size of bloodmeal determines the amount of virus initiating midgut infection, which directly determine the infection prevalence. In the infected mosquitoes, the smaller blood meal size lower the number of virus initiating midgut infection, which has previously been demonstrated to influence the level of persistent infection across the tissues (16,21). It is interesting to note that the comparative bloodmeal size between mosquito species and population has rarely been investigated despite its relevance to vector competence and vectorial capacity. In addition to vector competence, the differences in bloodmeal size may also determine the fecundity of female mosquitoes, thereby affecting mosquito population density. Our observations incite further investigation on the host and environmental factors determining bloodmeal size and the effect of bloodmeal size on mosquito biology.

The most concerning phenotype of *Ae. scutellaris* is its ability to support ZIKV and DENV replication and its low barrier for crossing tissue boundaries. This leads to high transmissibility of ZIKV and DENVs. In intrathoracic injection experiments, *Ae. scutellaris* showed comparable transmissibility to *Ae. aegypti* despite lower salivary gland ZIKV titers. Similarly, *Ae. scutellaris* had significantly higher transmissibility of DENV2 compared to *Ae. aegypti*.

While our study demonstrates the less permissive *Ae. albopictus* midgut for the establishment of midgut infection, there have been contradicting results regarding ZIKV and DENV establishment of midgut infection phenotype between *Ae. albopictus* and *Ae. aegypti*. Some studies found that *Ae. albopictus* had superior ability to support establishment of midgut infection to *Ae. aegypti* (22) while the others showed similar establishment of midgut infection (23,24) or the inferior midgut infection (25–28). These contradicting results demonstrate varying degrees of midgut permissiveness of different mosquito populations from the same species. Regardless of the midgut infectivity, most of these studies demonstrated similar or superior propagation of ZIKV and DENV in the tissues of *Ae. albopictus.* However, despite the high level of virus titers in the tissues, the salivary gland escape barrier is likely major limitation for ZIKV and DENV transmission by *Ae. albopictus*.

In addition to comparative vector competence phenotype between mosquito species, we were able to compare intra-host infection kinetics between ZIKV and DENV1-4 by controlling the bloodmeal or intrathoracic injection titers. We found that the kinetics of infection in the midgut, carcass and salivary glands were strikingly different among the viruses used in this study, suggesting tissue specific infection phenotype of different viruses. Specifically, ZIKV was the most infectious and transmissible virus compared to the DENVs. The virus was very efficient in establishment of midgut infection, propagation in secondary tissues and crossing intra-host infection boundaries, which eventually leads to the highest transmissibility. Despite a high transmissibility of ZIKV, the number of reported ZIKV cases in Thailand has been only a fraction of DENV cases suggesting that there might be other factors that limit ZIKV outbreak such as herd immunity in human populations (29). Another explanation is that ZIKV surveillance is much more limited than DENVs thus the reported number did not reflect a real number of infections. Among DENVs, the establishment of midgut infection was the highest in DENV3 followed by DENV1, DENV4 then DENV2. Our results were differ from a previous comprehensive intra-host infection kinetics by Novelo et al. (21), which showed that DENV1 and DENV2 were better than DENV3 and DENV4 in terms of the establishment of midgut infection. Interestingly, while DENV3 was very good at establishing midgut infection, the virus like has lower replication rate in the mosquitoes regardless of the vector species. In addition to DENV3, DENV4 also showed low replication rate in all mosquitoes, which was similar to what previously observed (21). The differences in the infection pattern observed between our study and previous studies highlighted the influence of vector and virus genotypes on the infection phenotypes (30,31).

It should be noted that, for practical reasons, our study conducted the infection experiments with only one strain per virus. Except for ZIKV that we used a local strain, the DENVs used in this study were the contemporary DENV panel freely available from BEI Resources (Cat no. NR-51131). The reason why we opted for these DENVs was to allow comparison between our results and other infection studies. Due to limited number of virus genotype used, it is possible that there may be specific interactions between virus and mosquito genotypes that influence infection outcome. Future works investigating the transmissibility of local mosquitoes and virus strains coupled with evaluation of vector population density in Zika and dengue endemic areas will provide a more comprehensive understanding on the role of each mosquito species in local virus transmission. Thailand offers a unique setting to conduct such future investigation due to an availability of weekly dengue case surveillance to district levels as well as research infrastructure.

The significance of our research is underscored by the potential of *Ae. scutellaris* in arbovirus circulation and outbreaks, especially considering changing climate patterns. Most studies on the geographical distribution of *Ae. scutellaris* date back over a decade (10,12,13). Given the high arbovirus transmission potential of *Ae. scutellaris* observed in our study, there is a crucial need for updated investigations into its geographical range and potential variations in arbovirus transmissibility. In addition to *Ae. scutellaris*, it is crucial to explore the vector competence of other members within the Scutellaris subgroup. While several studies have investigated the vector competence of *Ae. polynesiensis*, identifying its role as a vector for DENV, CHIKV, and ZIKV in the Pacific region (9,32), the potential involvement of other members in arbovirus transmission remains unestablished. Recognizing and understanding the role of this neglected vector species contributes not only to our understanding of disease transmission but also informs the development of effective vector control strategies.

## Acknowledgements

This work was supported by the Thailand Program Management Unit for Human Resources & Institutional Development, Research and Innovation (PMU-B), NXPO, grant number B05F640142 to NJ. The project was also supported by Mahidol university (Fundamental Fund: fiscal year 2023 by National Science Research and Innovation Fund (NSRF)) to PS. The following reagent was obtained through BEI Resources, NIAID, NIH: Contemporary Dengue Virus Panel, NR-51131. The Asian ZIKV SV0010/15 was obtained from the Armed Forces Research Institute of Medical Sciences (AFRIMS) and the Department of Disease Control, Ministry of Public Health through the Cluster Program Management Office, NSTDA. Illustrations of the figures were generated in Biorender (https://www.biorender.com).

## Declaration of Conflicting Interests

The author(s) declared no potential conflicts of interest with respect to the research, authorship, and/or publication of this article.

## Author contributions

Conceptualization: Chatpong Pethrak, Jutharat Pengon, Natapong Jupatanakul, and Patchara Sriwichai; Supervision: Natapong Jupatanakul, and Patchara Sriwichai.

Funding acquisition: Natapong Jupatanakul, and Patchara Sriwichai.

Investigation: Chatpong Pethrak, Jutharat Pengon, Saranya Thaiudomsup, Kittitat Suksirisawat, Thipruethai Phanitchat, Channarong Sartsanga, Tararat Jantra, Natapong Jupatanakul.

Formal analysis, and writing – original draft preparation: Chatpong Pethrak, Jutharat Pengon, Natapong Jupatanakul.

Resources: Jutharat Pengon, Chatpong Pethrak, Yudthana Samung, Anon Payakkapol, Songpol Eiamsam-ang, Thipruethai Phanitchat, Natapong Jupatanakul.

Visualization: Chatpong Pethrak, Natapong Jupatanakul

Writing – original draft, Chatpong Pethrak, Jutharat Pengon, and Natapong Jupatanakul.

Writing – review & editing, Chatpong Pethrak, Jutharat Pengon, Natapong Jupatanakul, and Patchara Sriwichai.

All authors read and approved the final manuscript.

## Supplementary Information

**Supplementary document S1**: Sequencing and Blastn results of Cytochrome Oxidase I (COI) from colonized *Aedes scutellaris*.

**Figure S1.**
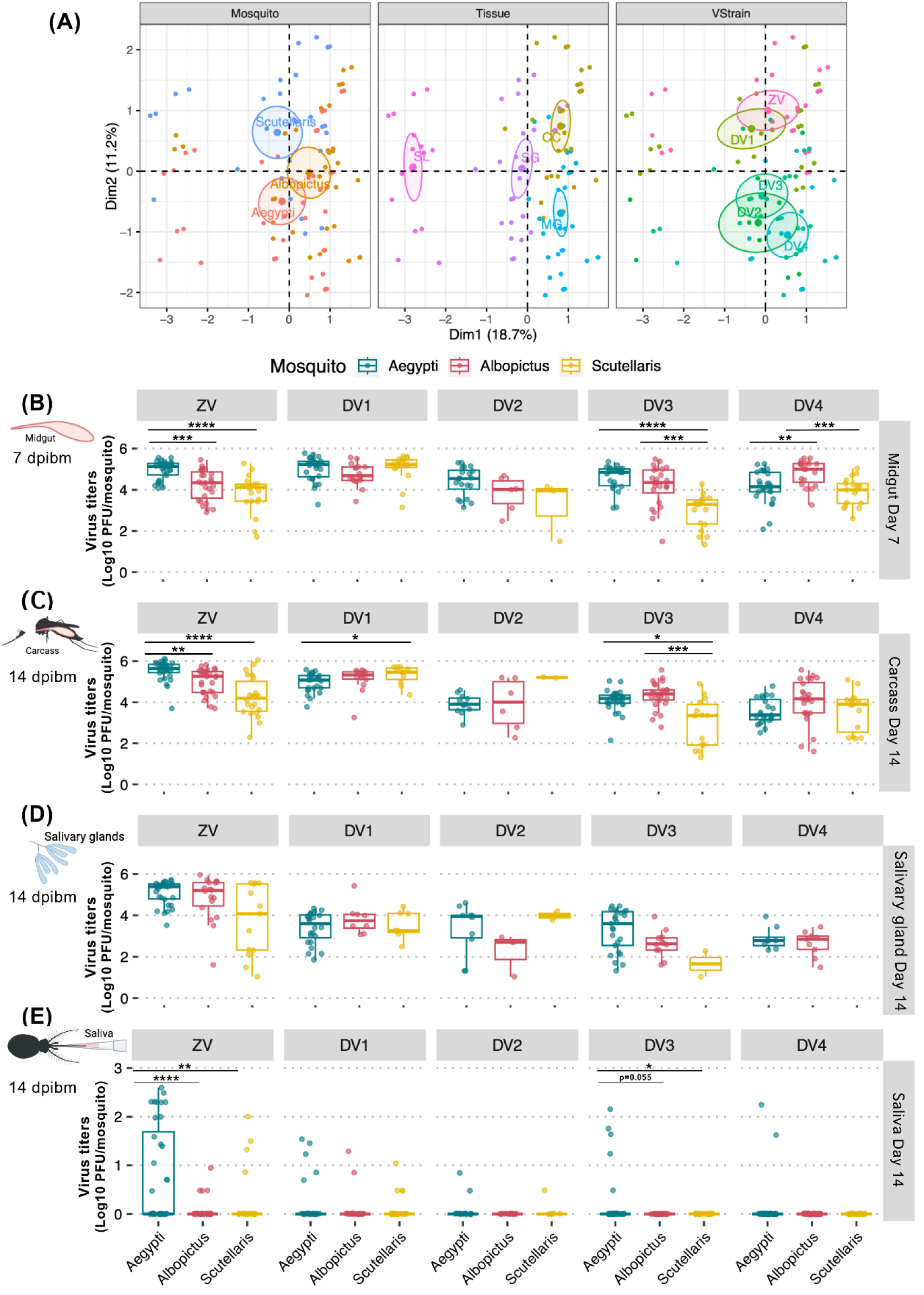
Comparison of ZIKV and DENV1-4 titers in different infection barriers of *Ae. aegypti*, *Ae. albopictus*, and *Ae. scutellaris*. (A) FAMD factor maps to visualize variance between median infection titers from each blood feeding experiment on the transformed coordinates. The information in the analysis included median infection titers, mosquito species (Mosquito), tissues type (Tissue), and virus (VStrain). The clustering is based on qualitative coordinates (mosquito species, tissue and virus), and each color represents individuals in a specific feature. Cluster distance represents the correlation among variables. The confidence ellipses indicate their potential relationships. Abbreviation: Dim, dimension. Box and scatter plot comparing infection titers of individual mosquito in the (B) midguts at 7 dpibm, (C) carcasses at 14 dpibm, (D) salivary glands at 14 dpibm, and (E) saliva at 14 dpibm grouping by mosquito species and virus. Each dot represents infection prevalence of from each blood feeding experiment. The feeding titers for each virus were 6.26-6.54 log_10_ PFU/mL for DENV1, 6.32-6.38 log_10_ PFU/mL for DENV2, 6.40-6.41 log_10_ PFU/mL for DENV3, 6.08-6.34 log_10_ PFU/mL for DENV4, and 6.32-6.56 log_10_ PFU/mL for ZIKV. Statistical analysis comparing infection level was conducted using Kruskal-Wallis followed by Dunn’s posthoc test in R. *: p<0.05, **: p<0.01.

**Figure S2.**
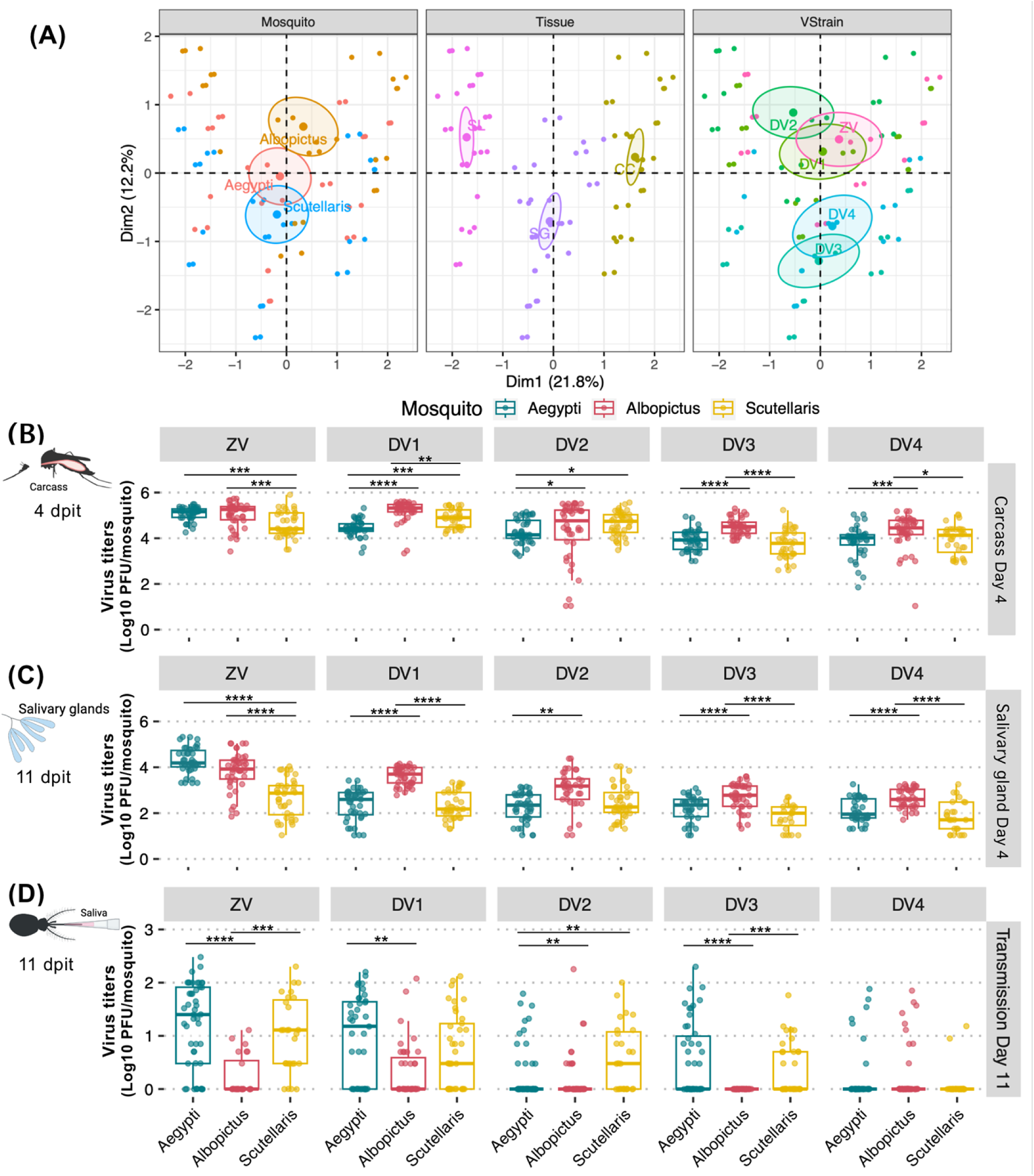
Comparison of ZIKV and DENV1-4 titers in different infection barriers of *Ae. aegypti*, *Ae. albopictus*, and *Ae. scutellaris* after intrathoracic injection. (A) FAMD factor maps to visualize variance between median infection titers from intrathoracic injection experiment on the transformed coordinates. The information in the analysis included median infection titers, mosquito species (Mosquito), tissues type (Tissue), and virus (VStrain). The clustering is based on qualitative coordinates (mosquito species, tissue and virus), and each color represents individuals in a specific feature. Cluster distance represents the correlation among variables. The confidence ellipses indicate their potential relationships. Abbreviation: Dim, dimension. Box and scatter plot comparing infection titers of individual mosquito in the (B) carcass at 4 dpit, (C) salivary glands at 4 dpit, and (D) saliva at 11 dpit, grouping by mosquito species or virus. Each dot represents infection prevalence of from each intrathoracic injection experiment. The titers for each virus were 6.23-6.40 log_10_ PFU/mL for DENV1, 6.11-6.34 log_10_ PFU/mL for DENV2, 5.85-6.40 log_10_ PFU/mL for DENV3, 6.11-6.41 log_10_ PFU/mL for DENV4, and 6.36-6.48 log_10_ PFU/mL for ZIKV. Statistical analysis comparing infection level was conducted using Kruskal-Wallis followed by Dunn’s posthoc test in R. *: p<0.05, **: p<0.01.

## Notes

### Competing Interest Statement

The authors have declared no competing interest.

